# Temporal emergence of functional dark matter in microbial responses to PFAS revealed by materials-based cultivation

**DOI:** 10.64898/2026.05.27.728134

**Authors:** Marta Velaz Martín, Kersten S. Rabe, Laura Meisch, John Vollmers, Anne-Kristin Kaster, Christof M. Niemeyer

**Affiliations:** Karlsruhe Institute of Technology (KIT), Institute for Biological Interfaces 1 (IBG-1), Biomolecular Micro- and Nanostructures, Hermann-von-Helmholtz-Platz 1, D-76344 Eggenstein-Leopoldshafen, Germany; Karlsruhe Institute of Technology (KIT), Institute for Biological Interfaces 5 (IBG-5), Microbial Genetics & Biotechnology, Hermann-von-Helmholtz-Platz 1, D-76344 Eggenstein-Leopoldshafen, Germany

## Abstract

Microbial responses to xenobiotic compounds are difficult to resolve due to environmental complexity and limited functional annotation. Here, we establish a materials-based cultivation framework using macroporous elastomeric silicone foams (MESIF) to capture microbial adaptation across environmental contexts and timescales. Using glyphosate as a model compound and per- and polyfluoroalkyl substances (PFAS) as a poorly understood class, we show that responses differ depending on the availability of established metabolic pathways. Glyphosate exposure induced rapid, pathway-specific functional enrichment with minimal taxonomic change. In contrast, PFAS exposure did not yield consistent taxonomic or annotation-based signals but instead produced responses that emerged over time and were primarily detectable at protein and genome-resolved levels. LC-MS analyses revealed transformation dynamics, including formation of shorter-chain products. These responses were not explained by known degrader taxa but involved uncharacterized proteins and microbial populations, highlighting the extent of functional “dark matter” in microbial responses to persistent contaminants.

## 1. Introduction

Microorganisms play a central role in maintaining environmental balance^[1–3]^. However, increasing human activity has led to the widespread release of persistent contaminants into natural ecosystems, posing significant challenges for environmental and human health^[4]^. These xenobiotics disperse into aquatic systems, contaminating freshwater and drinking water sources and ultimately reaching wastewater treatment plants (WWTPs)^[5]^.

Among these, glyphosate is one of the most widely used herbicides worldwide^[6]^. Beyond its effects on plants, glyphosate has been shown to impact microbial communities, including the gut microbiome^[7]^, as well as higher organisms^[8,9]^, with documented bioaccumulation and entry into the human body^[10–13]^. In contrast, per- and polyfluoroalkyl substances (PFAS) represent a more recalcitrant class of contaminants. These synthetic compounds are characterized by highly stable carbon-fluorine bonds^[14]^, resulting in extreme environmental persistence and resistance to degradation^[15,16]^. PFAS exhibit strong bioaccumulation potential and have been associated with adverse effects in higher organisms, including endocrine disruption in humans^[17, 18].^

While microbial processes are known to mediate the transformation of many environmental xenobiotics, mechanistic understanding varies widely across compound classes. Glyphosate degradation proceeds via well-characterized pathways, including cleavage of the C-P bond by the C-P lyase system and oxidation by glyphosate oxidoreductase (gox), yielding aminomethylphosphonic acid (AMPA)^[19–21]^. These pathways enable microorganisms to utilize glyphosate as a source of phosphorus and nitrogen. In contrast, the physicochemical properties of PFAS pose fundamental challenges to microbial metabolism, and their biological transformation remains poorly understood, limiting current strategies for their remediation^[22–24]^.

A major bottleneck in identifying microorganisms involved in xenobiotic transformation lies in their cultivation. Many environmentally relevant microbes remain uncultured due to unknown physiological requirements, slow growth, or dependence on community interactions^[25,26]^. Moreover, environmental systems such as WWTPs are inherently dynamic, with continuous mixing and periodic turnover of influent, which can obscure the detection of selective microbial responses over time. To address these limitations, cultivation-based enrichment platforms have emerged as powerful tools to capture microbial diversity directly from environmental samples^[27–29]^.

One such approach involves macroporous elastomeric silicone foams (MESIF) chips, which enable in situ cultivation under environmentally relevant conditions^[30]^. Previous work has shown that these platforms enrich distinct microbial communities compared to the native environment and respond to selective pressures such as glyphosate exposure^[30,31]^. However, whether such systems can be systematically applied to resolve microbial dynamics over time, particularly under conditions of contrasting environmental stability, and to link specific organisms and functions to xenobiotic exposure remains unclear.

Here, we establish a materials-based cultivation framework to selectively capture and characterize microbial responses to xenobiotic exposure. We use glyphosate as a model compound with well-defined degradation pathways and PFAS as a poorly understood compound class. MESIF chips were fabricated from a macroporous polydimethylsiloxane (PDMS) matrix generated by porogen leaching (Fig. 1a) and assembled into cultivation devices with an integrated reservoir enabling controlled and sustained exposure to xenobiotics (Fig. 1b,c). We implemented two complementary cultivation strategies: short-term in situ incubation in a WWTP to capture rapid responses under dynamic environmental conditions (Fig. 1d), and long-term laboratory mesocosm cultivation to resolve slower, emergent dynamics under more controlled conditions (Fig. 1e). All samples were processed using a unified workflow combining shotgun metagenomic sequencing with community-, protein-, and genome-resolved analyses (Fig. 1f). This integrated approach enables the identification of both rapid and temporally emerging microbial responses to xenobiotic exposure and reveals candidate organisms and functional traits, including previously uncharacterized protein space, associated with their transformation.

**Fig. 1.**
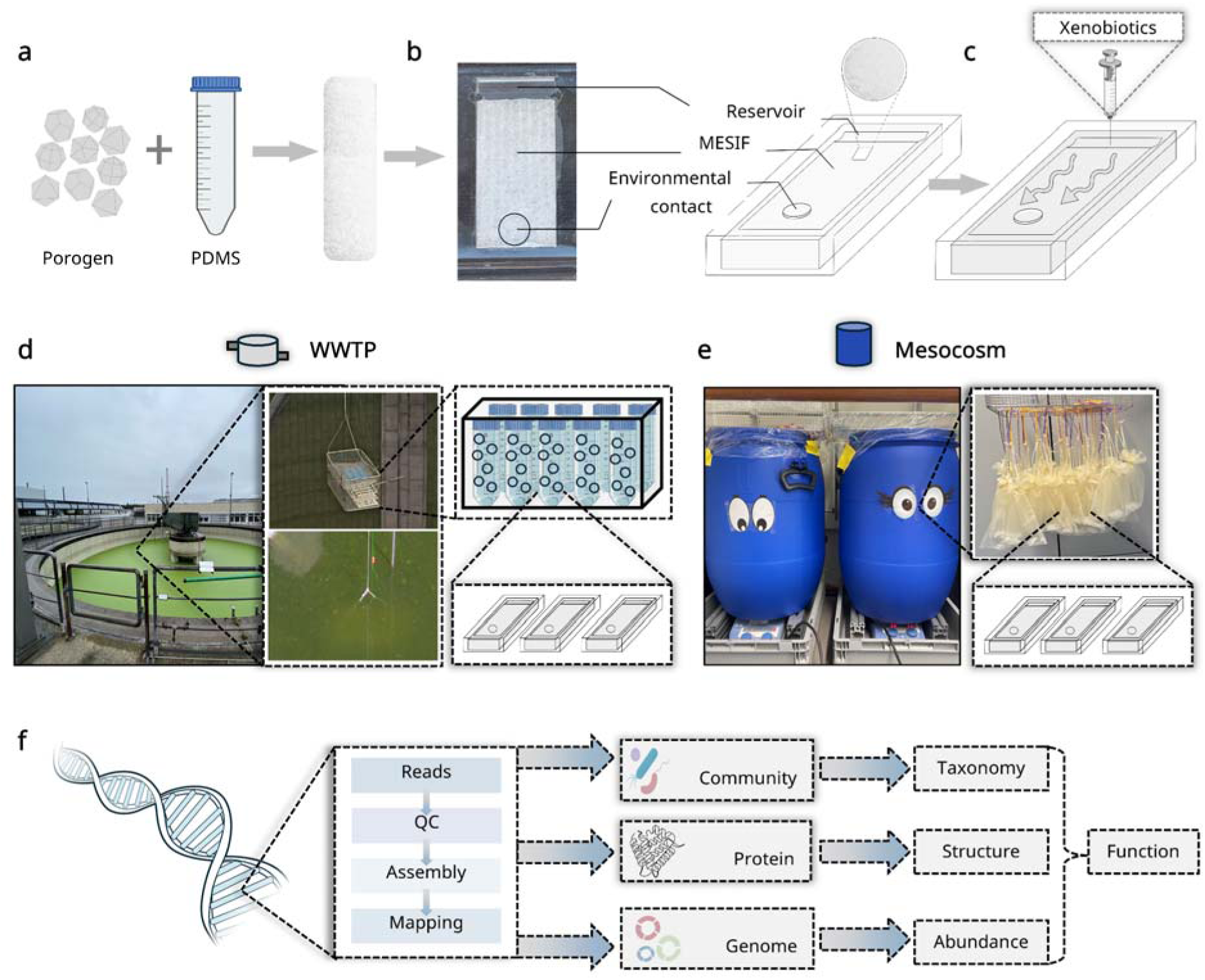
MESIF-based cultivation enables controlled and time-resolved enrichment of microbial communities under xenobiotic exposure. (a) MESIFs were fabricated by embedding porogens (salt crystals) into polydimethylsiloxane (PDMS), followed by leaching to generate a macroporous matrix. (b) MESIF chips consist of the porous matrix, a housing, and a perforated lid, allowing environmental exchange while maintaining a defined internal reservoir. (c) The reservoir enables controlled supplementation with basal media and xenobiotics. (d) For short-term in situ cultivation, MESIF chips were deployed in a wastewater treatment plant (WWTP) for 21 days. Chips were placed in perforated Falcon tubes, mounted in cages, and submerged in the treatment tank. (e) For long-term cultivation, MESIF chips were incubated in laboratory mesocosms for up to 100 days, placed in mesh bags and suspended in wastewater-filled tanks, using separate tanks for PFAS (left) and glyphosate (right) exposure (f) Following cultivation, samples were processed by DNA extraction and shotgun metagenomic sequencing, and analyzed at the community, protein, and genome-resolved levels to assess microbial taxonomy, abundance, structural features, and functional potential.

## 2. Results

To evaluate MESIF-based cultivation for resolving xenobiotic-associated microbial responses, we designed a multi-factorial framework spanning compound type, medium composition and cultivation timescale. MESIF chips were supplemented with xenobiotics either through the integrated reservoir or by adsorption to the matrix, depending on compound properties, and incubated in defined minimal medium (M9) or wastewater-derived medium (N). Cultivation was performed as short-term in situ incubation in a WWTP and as long-term laboratory mesocosm incubation to capture rapid and temporally evolving responses. We first used glyphosate as a benchmark compound with well-characterized degradation pathways, before extending the framework to PFAS, for which microbial transformation remains poorly understood.

### 2.1. Platform validation with glyphosate reveals functional adaptation

To evaluate selective enrichment of xenobiotic-responsive microbiomes, we used glyphosate as a model compound. MESIF chips were supplemented with glyphosate or left untreated and incubated in defined minimal medium (M9) or wastewater-derived medium (N). Given the high solubility of glyphosate, compound retention was assessed using the fluorescent tracer sodium fluorescein. Only minimal diffusion into the surrounding medium (∼0.5%) was observed, indicating sustained retention within the matrix (Supplementary Fig. S1).

To establish a baseline, we compared microbial communities recovered from MESIF chips without xenobiotic supplementation (M9 and N) to those in the surrounding habitat. MESIF cultivation resulted in pronounced taxonomic and functional shifts, including increased recovery of low-abundance taxa and enhanced genome reconstruction (Extended Data Fig. 1). Taxonomic composition at the phylum level was assessed based on SSU and LSU rRNA genes (Extended Data Fig. 2). Community profiles differed across media, systems and time. In the WWTP, MESIF communities were dominated by *Pseudomonadota* and *Campylobacterota*, whereas habitat communities remained more evenly distributed. In mesocosms, MESIF communities showed stronger temporal turnover, with early dominance of *Pseudomonadota* followed by decline at later time points, particularly in N-derived samples (Extended Data Fig. 2b,c). Together, these results indicate that MESIF enrichment is context dependent and shaped by medium, incubation setting and time^[30,31]^.

Having established baseline differences between MESIF-derived communities and the surrounding habitat, subsequent analyses focused on xenobiotic-supplemented MESIF chips (M9-G and N-G) relative to their respective controls (M9 and N) within each system (WWTP and mesocosm), thereby minimizing cultivation-derived biases. At the taxonomic level, glyphosate supplementation resulted in only minor changes in overall community composition, with limited shifts at both phylum and genus levels, indicating that taxonomic restructuring is not the primary driver of glyphosate-associated responses (Extended Data Fig. 3, Supplementary Fig. S2).

We next assessed functional responses to glyphosate supplementation. Genes associated with glyphosate degradation, including C-P lyase systems, AMPA-related pathways, phosphonate transporters and sarcosine metabolism, were consistently enriched (Fig. 2a,b), indicating selection on metabolic potential rather than taxonomic identity. Enrichment was more pronounced in M9, whereas N showed a weaker signal, consistent with differences in nutrient availability.

**Fig. 2.**
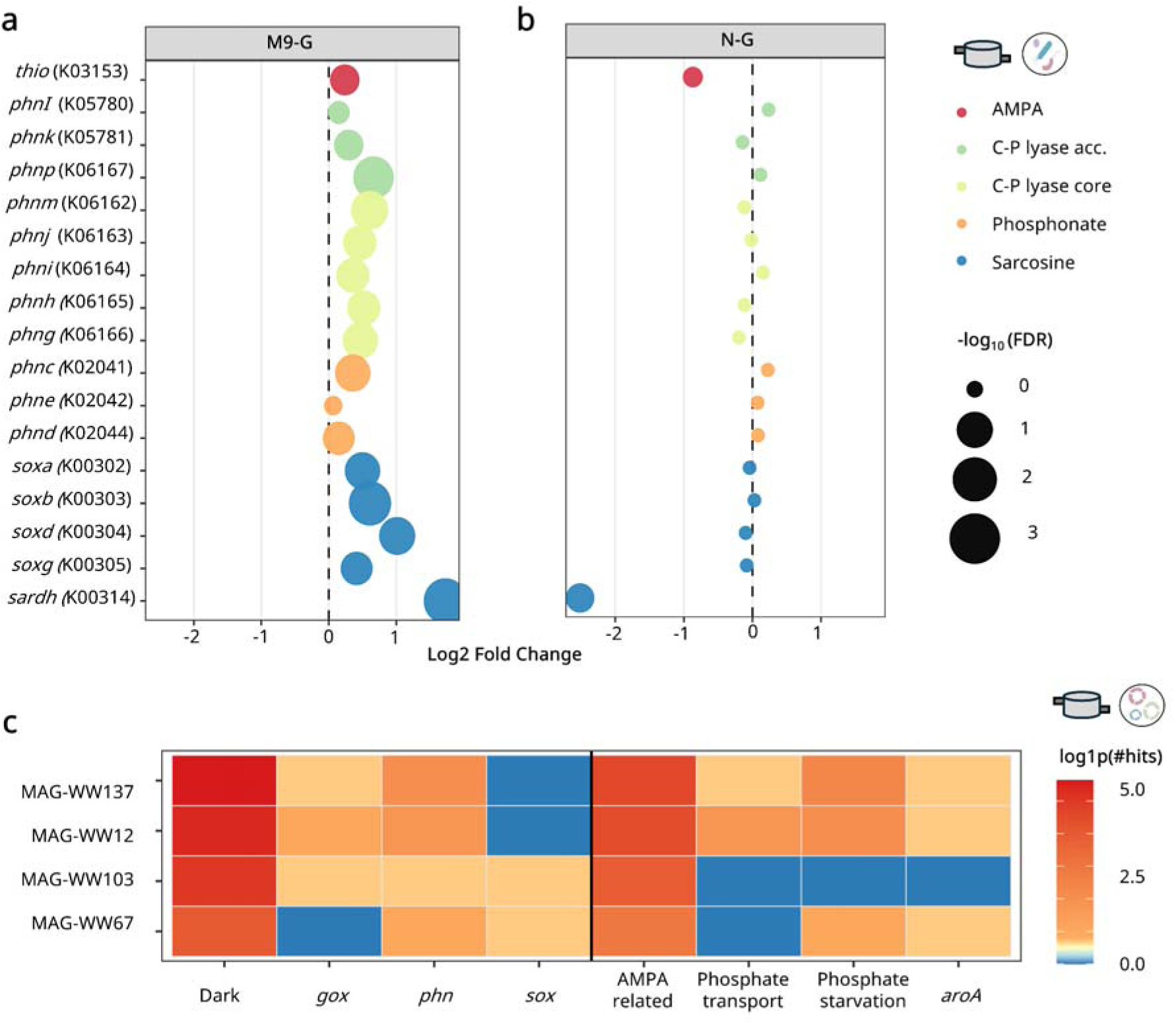
Glyphosate supplementation drives pathway-specific functional enrichment during short-term WWTP cultivation. (a,b) Functional response of microbial communities to glyphosate supplementation in (a) M9 and (b) native wastewater-derived medium (N). Log_2_ fold changes are shown relative to the corresponding control without glyphosate supplementation. Each point represents a gene associated with key glyphosate-related pathways, including AMPA metabolism (red), the C-P lyase system (core and accessory components; green and yellow), phosphonate transport (orange), and sarcosine metabolism (blue). Point size reflects statistical significance (−log_10_ FDR). (c) Functional profiling of metagenome-assembled genomes (MAGs) uniquely detected in glyphosate-supplemented samples. The heatmap shows the abundance of pathway-related gene hits per MAG, with color intensity indicating the number of detected hits.

PFAM-based analysis further revealed enrichment of mobile genetic element-associated domains under glyphosate supplementation (Extended Data Fig. 4a,b), suggesting elevated stress responses and potential horizontal gene transfer, consistent with the dissemination of glyphosate-degrading genes across microbial communities as previously reported^[32]^.

Next, we performed genome-resolved analyses to identify metagenome-assembled genomes (MAGs) associated with glyphosate supplementation. In total, 174 MAGs were reconstructed across all WWTP samples, including both control and glyphosate-supplemented conditions (Supplementary Table S1). Among these, four were exclusively detected under glyphosate supplementation (Extended Data Fig. 4c) and encoded key phosphonate degradation pathways, including C-P lyase systems and associated transporters, indicating their potential involvement in glyphosate transformation (Fig. 2c).

A similar pattern was observed in the mesocosm, where 388 MAGs were reconstructed (Supplementary Table S2), including 13 uniquely associated with glyphosate supplementation (Supplementary Fig. S3). In contrast to the WWTP, this increase in genome-resolved diversity was not accompanied by consistent functional enrichment (Supplementary Fig. S4). Instead, community variation was primarily driven by temporal dynamics, with prolonged incubation attenuating initial selection effects and reducing differences between supplemented and control communities (Supplementary Figs. S5-S6). These results suggest that responses to compounds with well-defined degradation pathways can be rapidly established but become less distinct over time.

To validate the enrichment approach, we assessed the presence of known glyphosate degraders by mapping reference genomes to metagenomic datasets (Supplementary Table S3). In the WWTP, only a weak signal from a single degrader was detected, whereas in the mesocosm signals were stronger but limited to two of six members (Supplementary Fig. S7a,b). Detection in control samples indicates that these organisms are part of the background community rather than exclusively associated with glyphosate supplementation. A diffusion assay using fluorescein revealed minimal release into the surrounding medium, indicating sustained retention within the MESIF matrix (Supplementary Fig. S1). This suggests localized exposure conditions that may promote distributed metabolic activity across multiple taxa rather than dominance of specialist degraders. Consistent with this, functional enrichment of glyphosate-associated pathways was observed despite limited detection of canonical degraders, highlighting contributions from low-abundance or uncharacterized taxa.

Together, these results indicate that functional responses to glyphosate supplementation are primarily driven by well-characterized metabolic pathways and can emerge without pronounced shifts in community composition, reflecting a distributed metabolic potential across diverse and often low-abundance taxa. In contrast to glyphosate, where degradation mechanisms are well defined and rapidly activated^[10,20, 32–37]^, PFAS remain poorly understood with respect to their biological transformation^[16,38]^. We therefore used PFAS as a second use case to assess whether microbial responses extend beyond known pathway-driven mechanisms and involve previously uncharacterized functional space.

### 2.2. Protein clustering uncovers PFAS-associated dark functional space

To extend the MESIF platform to xenobiotics lacking well-defined degradation pathways, we investigated PFAS using perfluorooctanoic acid (PFOA) and perfluorooctanesulfonic acid (PFOS). Due to their limited solubility, both compounds were introduced via physisorption onto the MESIF matrix. LC-MS measurements showed minimal release, indicating stable retention during cultivation (Supplementary Fig. S8,S9). MESIF chips were cultivated in WWTP and mesocosm systems, followed by taxonomic and functional profiling across control (M9, N) and PFAS-supplemented conditions.

In contrast to glyphosate, PFAS exposure did not result in consistent taxonomic enrichment across conditions, with only minor genus-level shifts observed (Extended Data Figs. 5-6; Supplementary Figs. S10-S11). Community variation was primarily driven by medium rather than PFAS exposure. A limited number of genera showed reproducible increases, including *Pseudomonas* in the WWTP and *Aeromonas* in the mesocosm. Given the association of *Pseudomonas* with xenobiotic transformation^[16,39, 40]^, this may indicate PFAS-associated responses, although patterns remained taxonomically diffuse. Notably, taxonomic structure increased during prolonged mesocosm incubation but remained insufficient to explain PFAS-associated functional responses.

Functional changes in annotated PFAS-associated categories were weak and inconsistent (Extended Data Fig. 7a,b), indicating that PFAS-associated functional potential is not captured by annotation-based approaches and may instead reside within the “functional dark matter” of the community, i.e. proteins lacking functional annotation^[41,42]^. To address this, we applied protein-level clustering across all conditions, including both annotated and unannotated sequences. Clusters present across all conditions were excluded to reduce background redundancy and enrich for condition-associated signals. Uncharacterized proteins from these clusters were selected for downstream analyses.

Protein-level clustering revealed treatment-associated groups, including clusters shared between PFOA- and PFOS-exposed samples in the WWTP (Fig. 3a,b). However, many of these clusters were also detected in control samples, indicating partial enrichment rather than strict PFAS specificity (Fig. 3c,d). Notably, this pattern shifted in the mesocosm, where the proportion of treatment-specific proteins increased compared to the WWTP incubation (Fig. 3e,f; Extended Data Fig. 7c,d), suggesting that prolonged incubation enhances the resolution of PFAS-associated protein modules and enables the emergence of condition-associated functional patterns. Temporal profiling of these proteins further showed that they were predominantly detected in PFAS-exposed samples and became more consistently resolved over time (Supplementary Figs. S12-S13), supporting their classification as condition-associated candidates.

**Fig. 3.**
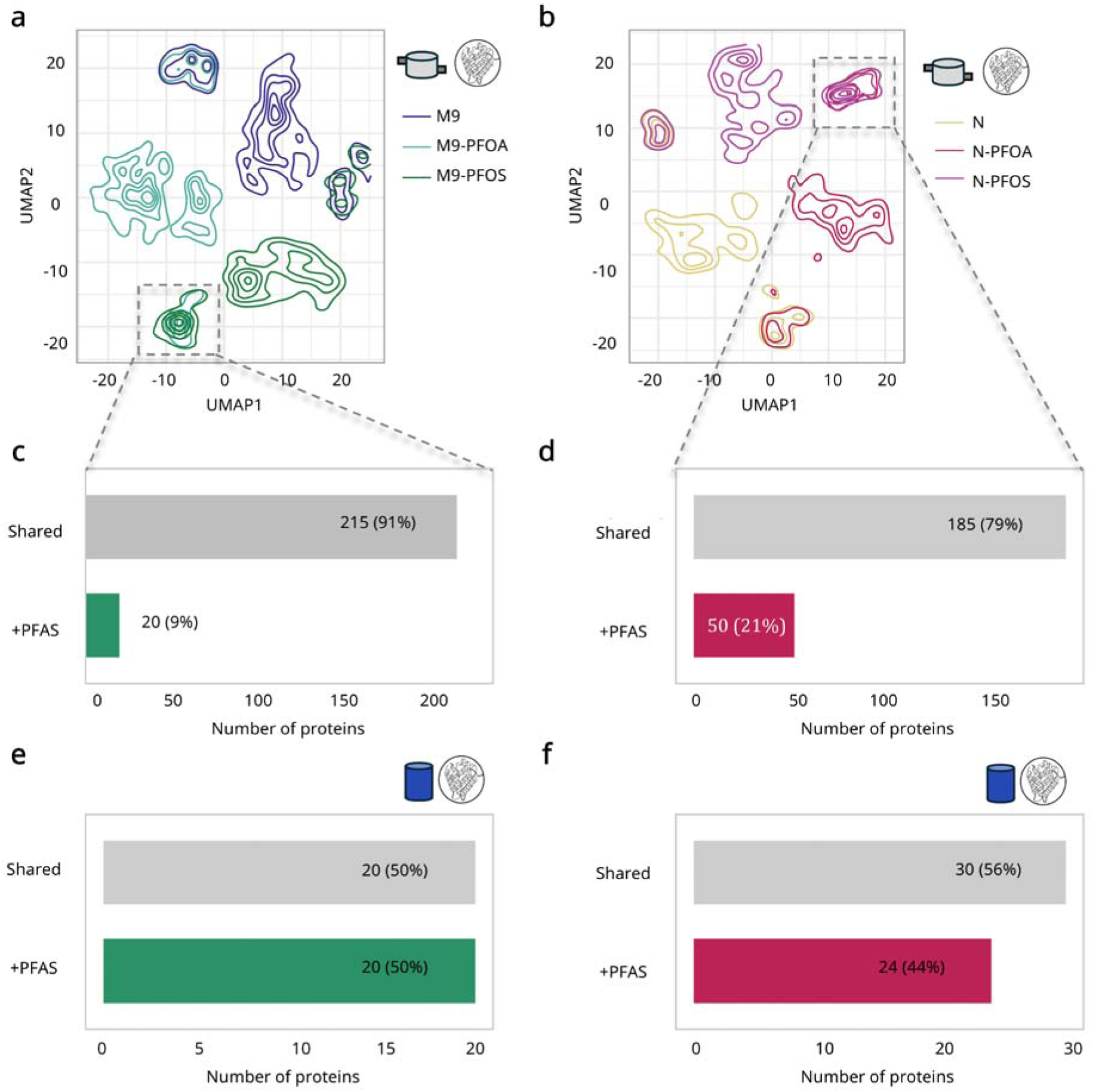
Protein-level clustering reveals treatment-associated modules and increased specificity during mesocosm cultivation. (a,b) UMAP projection of proteins from short-term WWTP cultivation after filtering to remove broadly shared proteins. Only proteins present in at most n-1 conditions were retained, enriching for condition-dependent signals. Density contours indicate the distribution across treatments (control, PFOA, PFOS) for (a) M9 and (b) native (N) communities, highlighting both partially shared and treatment-associated regions of functional space. (c,d) Quantification of PFAS-associated proteins in WWTP samples. Bars indicate the number and proportion of proteins detected in both PFOA- and PFOS-treated samples (+PFAS) compared to those also present in controls (Shared) for (c) M9 and (d) N. (e,f) Equivalent analysis for mesocosm samples, showing an increased proportion of treatment-associated proteins compared to WWTP for (e) M9 and (f) N.

To further characterize PFAS-associated uncharacterized proteins, we predicted structures using AlphaFold2 combined with Foldseek-based annotation (Supplementary Tables S4-S7). Across all treatments, at least 50% of candidates remained structurally uncharacterized, supporting the presence of functional dark matter (Extended Data Fig. 7e). Structurally dark proteins, defined here as sequences without detectable Foldseek hits, were preferentially recovered in native (N) medium-derived samples but not in M9 (Supplementary Table S8), indicating that medium composition influences the recovery of uncharacterized proteins. Together, these results suggest that PFAS-associated functional potential may reside within previously uncharacterized protein modules. However, it remains unclear whether these signals reflect biodegradation or tolerance-based responses, and protein-level clustering alone does not resolve which community members encode these functions or are specifically enriched under PFAS exposure.

To address this, we performed genome-resolved analyses of metagenome-assembled genomes (MAGs) uniquely detected in PFAS-treated samples (PFOA, PFOS, or both), assessing their contribution to the identified clusters and their potential involvement in PFAS-associated processes.

### 2.3. Genome-resolved analysis identifies PFAS-associated populations and functions

Building on the identification of PFAS-associated protein clusters, we performed genome-resolved analyses to identify metagenome-assembled genomes (MAGs) preferentially detected in PFAS-treated samples. In total, 14 such MAGs were identified in the WWTP (Fig. 4a) and 34 in the mesocosm (Supplementary Fig. S14). We functionally annotated these genomes to assess their potential contribution to PFAS-associated processes^[22]^.

**Fig. 4.**
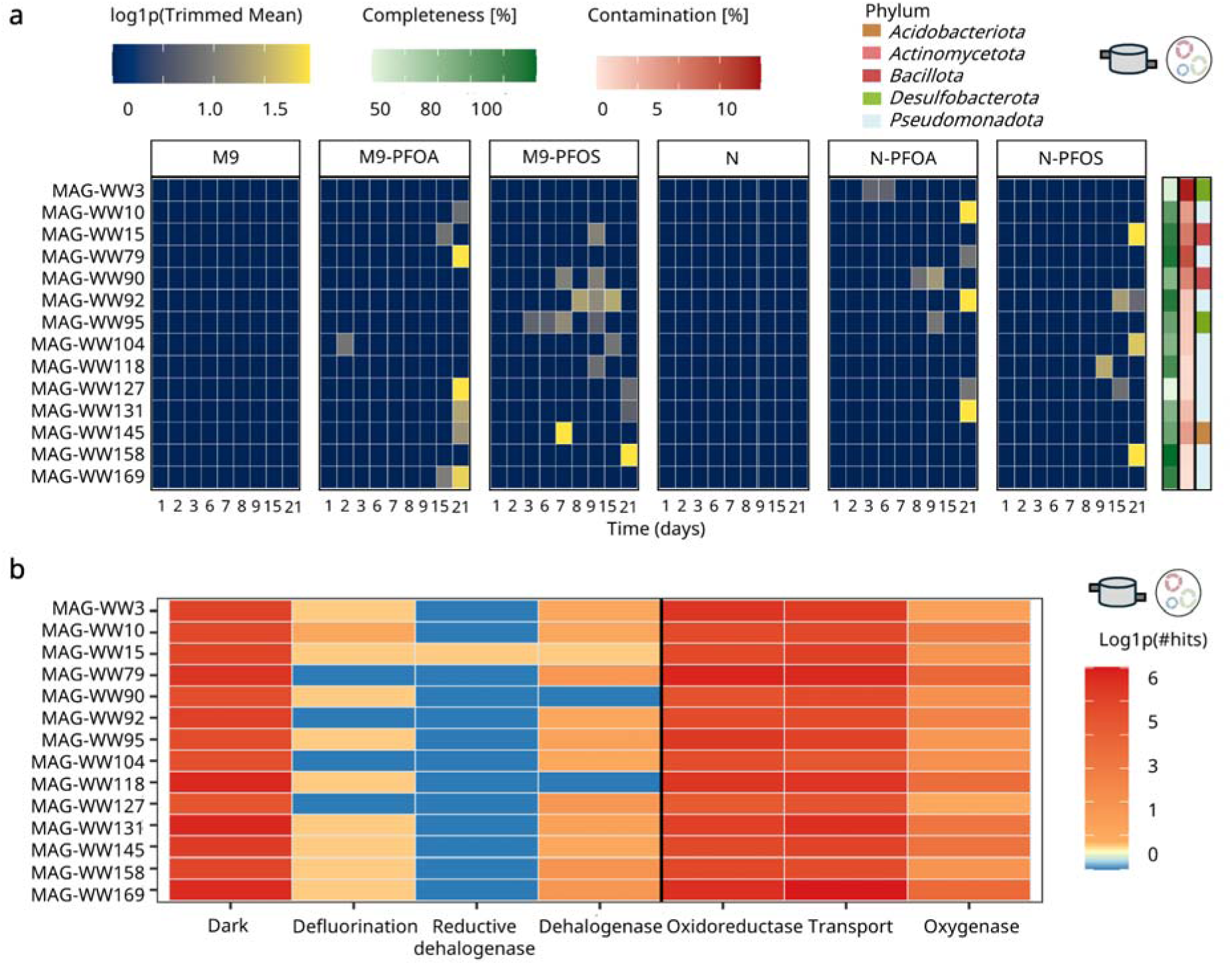
Genome-resolved analysis identifies PFAS-associated MAGs and their functional potential. (a) Temporal detection of metagenome-assembled genomes (MAGs) preferentially detected in PFAS-treated samples during short-term WWTP cultivation. Rows represent individual MAGs and columns correspond to sampling timepoints, with color intensity indicating relative abundance. Bars on the right indicate genome completeness and contamination, and phylum-level classification is shown alongside. (b) Functional annotation of PFAS-associated MAGs based on gene hits related to processes potentially relevant in PFAS-impacted systems, including dehalogenation, redox activity, and stress response. The heatmap shows the number of detected genes per functional category.

PFAS-associated MAGs were more frequently detected at later timepoints in both WWTP and mesocosm cultivation (Fig. 4a; Supplementary Fig. S14), with similar temporal patterns observed across independent systems, suggesting that condition-associated populations become more apparent as incubation progresses, consistent with the temporal emergence of PFAS-associated functional signals. Functional annotation revealed genes linked to processes potentially relevant in PFAS-impacted systems, including dehalogenases, oxidoreductases, transport systems, as well as a substantial fraction of uncharacterized proteins (Fig. 4b; Supplementary Fig. S15).

Notably, one MAG reconstructed from the WWTP encoded a reductive dehalogenase-like protein (Fig. 4b; MAG-WW15). Reductive dehalogenases are key enzymes mediating organohalide respiration^[43]^, catalyzing the reductive cleavage of carbon-halogen bonds under anaerobic conditions. Although PFAS are highly recalcitrant due to the strength of the C-F bond^[22]^, dehalogenase-mediated mechanisms have been proposed as potential biological routes for their transformation^[43–47]^, suggesting a possible role of this MAG in PFAS-associated processes.

Inspection of the genomic neighborhood revealed a conserved locus organization, in which the rdh-like gene is flanked by multiple uncharacterized (dark) proteins, a membrane-associated MBOAT protein, and a downstream chemotaxis-related sensor (Extended Data Fig. 8a). Given recent evidence that PFAS transformation intermediates can be incorporated into bacterial membrane lipids^[48]^, the presence of a membrane-associated enzyme may indicate potential involvement in membrane-related adaptation or interaction processes. Finally, phylogenetic placement of this MAG showed that it clusters within the genus *Fusibacter* (Extended Data Fig. 8b), suggesting that it represents a previously uncharacterized lineage within this group.

Mapping of reference genomes from previously reported PFAS-associated degraders revealed detectable signals in PFAS-exposed samples, with generally higher abundance in the mesocosm compared to WWTP conditions (Supplementary Fig. S16). However, these signals were inconsistent across timepoints and treatments, indicating that known degraders do not dominate PFAS-associated community responses in this system. This observation is consistent with the emergence of distinct PFAS-associated MAGs and protein clusters identified above. This further supports the notion that PFAS-associated functional responses are not primarily driven by previously characterized degrader taxa.

Integration of protein- and genome-level data showed partial overlap between MAG-derived and reference proteins in WWTP samples, whereas mesocosm samples exhibited more distinct clustering, consistent with increased resolution of condition-associated protein modules (Supplementary Fig. S17). Unassigned proteins remained broadly distributed, highlighting functional diversity beyond characterized protein families.

To further assess PFAS-associated processes, we monitored transformation products during mesocosm cultivation using LC-MS (Fig. 5a). PFOA concentrations decreased over time in MESIF samples and were accompanied by the formation of shorter-chain products (C4-C8), whereas PFOS remained largely stable with limited byproduct formation, indicating compound-specific transformation dynamics (Fig. 5b,c).

**Fig. 5.**
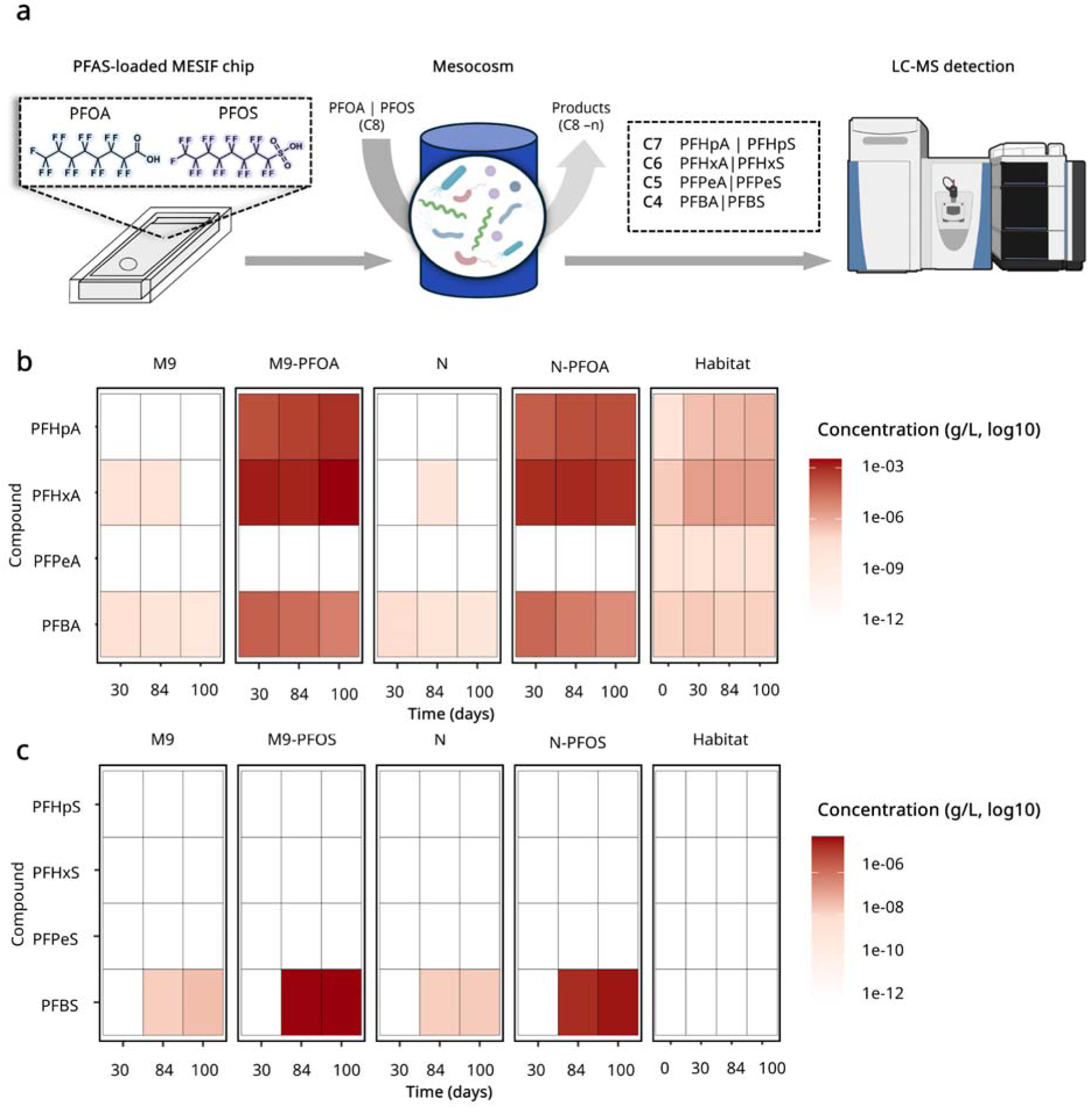
PFAS transformation dynamics during mesocosm cultivation in MESIF chips. (a) Schematic overview of the experimental setup. PFAS (PFOA, 18 mM; PFOS, 14.8 mM; initial loading concentration 7.4 g L⁻¹) were physisorbed onto MESIF chips and cultivated in mesocosms, followed by LC-MS monitoring of parent compounds and transformation products (C4-C8). (b,c) Heatmaps of (b) PFOA-derived and (c) PFOS-derived transformation products across time points and conditions. PFOA concentrations decreased over time and were accompanied by the formation of shorter-chain products, whereas PFOS remained largely stable with limited detectable byproducts.

### 2.4. Cross-system comparison reveals conserved PFAS-associated signatures

Finally, we investigated the presence and potential correlation of PFAS-associated signals between WWTP and mesocosm systems. To this end, we compared the most enriched and depleted genera across both systems relative to their respective controls to identify taxa consistently associated with PFAS exposure.

Overall, taxonomic responses differed substantially between experimental systems (Supplementary Fig. S18). Despite this variability, a subset of genera was reproducibly enriched across PFAS treatments and media, including *Pseudomonas*, a genus previously associated with xenobiotic transformation^[49]^, as well as sulfate-reducing bacteria such as *Desulfovibrio* and *Desulfomicrobium* (Supplementary Fig. S19). These taxa are linked to anaerobic reductive metabolism, including dehalogenation-related processes relevant for the transformation of highly halogenated compounds^[45]^. Together, these observations indicate a limited but reproducible taxonomic response that becomes more pronounced over time. Consistent with this, the magnitude of enrichment was generally higher in mesocosm samples compared to WWTP-derived samples, suggesting that prolonged incubation enhances the resolution of PFAS-associated taxonomic patterns.

We next examined protein clusters shared across both systems. No consistent clustering patterns were observed, indicating the absence of a conserved global protein-level signature (Extended Data Fig. 9a,b). However, 64 proteins were consistently detected across both systems (Extended Data Fig. 9a,b; black triangles), including 11 sequence-dark proteins lacking functional annotation (Extended Data Fig. 9c; Supplementary Table S10). To further characterize these candidates, we used AlphaFold2-predicted structures in combination with Foldseek-based annotations. To assess their potential for PFAS interaction, all structures were analyzed using pocket prediction (fpocket)^[50]^ followed by docking simulations (gnina)^[51]^. A control panel of known PFAS-binding and non-binding proteins clearly distinguished positive and negative controls (Supplementary Table S11), supporting the robustness of the approach. Binding affinities across candidates varied (Supplementary Table S12), with five proteins showing scores comparable to positive controls (Supplementary Table S13). Structural analysis of these candidates revealed diverse predicted binding pockets (Supplementary Fig. S20). While these results do not demonstrate catalytic activity, they suggest that these proteins have the capacity to interact with PFAS compounds.

Together, these findings indicate that PFAS-associated functional potential extends beyond annotated pathways and may reside in previously uncharacterized proteins, highlighting candidates for future experimental validation.

## 3. Discussion

We established a materials-based cultivation framework to resolve microbial responses to xenobiotic exposure across environmental contexts and timescales. Using glyphosate as a model compound and PFAS as a poorly understood class, we show that MESIF-based cultivation captures distinct modes of microbial adaptation that depend on compound properties and temporal dynamics. The porous matrix enables surface-associated growth under sustained local exposure, facilitating recovery of slow-growing and environmentally adapted microorganisms. Across systems, responses were not consistently reflected at the taxonomic level but instead emerged at functional and genome-resolved levels.

For glyphosate, microbial responses were characterized by rapid and pathway-specific functional enrichment, consistent with the activation of well-established degradation mechanisms. Despite only minor shifts in community composition, genes associated with phosphonate metabolism, including C-P lyase systems and AMPA-related pathways, were consistently enriched, indicating that selection primarily acted on metabolic potential rather than taxonomic identity. These responses were most pronounced under nutrient-limited conditions, where glyphosate served as a key resource, highlighting the importance of environmental context in shaping functional adaptation. In contrast, prolonged incubation in the mesocosm led to a partial attenuation of these signals, suggesting that, for compounds with defined metabolic pathways, selective responses can be rapidly established but may become less distinct as communities equilibrate over time.

In contrast to glyphosate, PFAS exposure did not induce strong or consistent responses at the taxonomic or annotation-based level. Instead, PFAS-associated signals emerged over time and were primarily detectable through protein-level and genome-resolved analyses. Protein clustering revealed condition-associated modules that became more distinct during prolonged mesocosm cultivation, while genome-resolved analyses identified MAGs that increased in prevalence over time. Consistent with these observations, LC-MS analyses indicated compound-specific transformation dynamics, with changes in parent compounds and formation of shorter-chain products. Differences between PFAS types likely reflect variation in physicochemical properties and bioavailability, influencing microbial accessibility and transformation potential.

These findings point to a fundamental difference in how microbial communities respond to xenobiotics depending on the availability of established metabolic pathways. While glyphosate responses were driven by the activation of known and conserved biochemical routes, PFAS-associated responses appear to involve a broader and less well-defined functional repertoire. Notably, known PFAS-associated degrader taxa were detected only at low abundance and did not dominate community responses, indicating that the observed signals are not primarily explained by previously characterized organisms. Instead, both protein-level and genome-resolved analyses consistently pointed to the involvement of previously uncharacterized proteins and microbial populations.

A key insight from this study is the importance of temporal dynamics in resolving microbial responses to recalcitrant compounds. While short-term in situ cultivation in the WWTP captured rapid responses under highly dynamic conditions, longer-term mesocosm cultivation enabled the emergence of more structured and reproducible patterns. This is consistent with the inherent differences between these systems, where continuous mixing and turnover in WWTP environments may obscure selective processes, whereas more stable conditions in mesocosms allow gradual enrichment of responsive populations. Importantly, even under prolonged incubation, taxonomic signals remained only partially informative, reinforcing the notion that functional and genome-resolved analyses are required to capture the full extent of PFAS-associated microbial responses.

The identification of uncharacterized protein clusters and MAG-associated functions highlights the extent of functional “dark matter” involved in PFAS-associated processes. Structural analyses further indicated that a subset of these proteins may possess the capacity to interact with PFAS compounds, although their precise biochemical roles remain to be determined. Together, these findings suggest that microbial responses to PFAS are likely mediated by distributed and previously unrecognized functional traits rather than by a small number of specialized degraders.

Several limitations should be considered. While MESIF-based cultivation enables sustained local exposure to xenobiotics and facilitates the recovery of responsive communities, it represents a semi-controlled system that may not fully capture all environmental processes. In addition, although preliminary analyses suggest temporal changes in PFAS concentrations and potential transformation products, further targeted chemical analyses are required to directly link observed microbial responses to specific transformation pathways. Finally, functional predictions based on sequence clustering and structural modeling provide valuable hypotheses but require experimental validation to confirm biochemical activity.

In summary, MESIF-based cultivation provides a framework to resolve microbial responses to xenobiotic exposure across environmental and temporal gradients. By integrating community, protein-level and genome-resolved analyses, this approach reveals functional adaptation processes that remain inaccessible to annotation-based methods. These findings advance our understanding of microbial interactions with persistent contaminants and highlight the importance of integrating cultivation-based enrichment with multi-level molecular analyses to uncover previously hidden functional potential.

## 4. Methods

### 4.1. MESIF cultivation platform

#### 4.1.1. MESIF chip preparation

Macroporous elastomeric silicone foams (MESIF) were fabricated as previously described.^[30]^ In brief, polydimethylsiloxane (PDMS; SYLGARD® 184) was mixed with curing agent at a 10:1 (w/w) ratio, degassed under vacuum, and cast into polymethylmethacrylate (PMMA) molds containing sieved salt crystals (500-707 µm) that served as porogens. After centrifugation to facilitate porogen embedding, the PDMS matrix was cured at 70 °C. The solidified PDMS-salt composites were removed from the molds and salt was leached by repeated washing in warm water to generate a macroporous structure (Fig. 1a). The resulting foams were dried and cut to the defined dimensions (18 x 10 x 3 mm). PDMS housings (cages and lids) were fabricated separately using PMMA molds and cured at 70 °C. Lids were perforated to allow direct medium contact with the porous matrix. MESIF foams and PDMS housings were subsequently treated with oxygen plasma to activate bonding surfaces and assembled into complete MESIF chips, followed by thermal curing under mechanical pressure to ensure stable sealing (Fig. 1b).

#### 4.1.2. Cultivation media

In this study, two cultivation media were used: a defined mineral medium (M9) and a complex environmental medium, referred to hereafter as native medium (N). M9 consisted of 1× M9 salts supplemented with 1 mM MgSO₄, 0.3 mM CaCl₂, 1 mg l⁻¹ biotin, 1 mg l⁻¹ thiamine and 1× trace element solution. The M9 salts were prepared from a 10× stock solution (75.2 g l⁻¹ Na₂HPO₄·2H₂O, 30 g l⁻¹ KH₂PO₄, 5 g l⁻¹ NaCl and 5 g l⁻¹ NH₄Cl), and the trace element solution was prepared from a 100× stock solution (5 g l⁻¹ EDTA, 0.83 g l⁻¹ FeCl₃·6H₂O, 84 mg l⁻¹ ZnCl₂, 13 mg l⁻¹ CuCl₂·2H₂O, 10 mg l⁻¹ CoCl₂·2H₂O, 10 mg l⁻¹ H₃BO₃ and 1.6 mg l⁻¹ MnCl₂·4H₂O). The native medium consisted of wastewater that was autoclaved at 120 °C for 20 min and subsequently sterile filtered through a 0.2 µm membrane before use. Given its origin, the N medium was expected to contain a higher and more chemically diverse pool of organic carbon than M9, in which carbon availability was limited to the supplied xenobiotic compounds.

#### 4.1.3. Supplementation of xenobiotics and medium

Xenobiotic supplementation was performed using two different strategies depending on the physicochemical properties of each compound, with all compounds delivered via the reservoir (Fig. 1c). The water-soluble herbicide glyphosate (Roundup™, Bayer, German) was directly added to the cultivation medium at a final concentration of 40 g L⁻¹ (0.24 M) at the time of inoculation (Supplementary Fig. S22a). This concentration was selected to ensure strong selective pressure and therefore exceeds environmentally relevant levels.^[12]^

In contrast, per- and polyfluoroalkyl substances (PFAS) were obtained from Merck and introduced via physisorption onto the MESIF matrix due to their limited aqueous solubility. Stock solutions of perfluorooctanoic acid (PFOA) and perfluorooctanesulfonic acid (PFOS) (7.4 g L⁻¹ in LC/MS-grade methanol) corresponding to 18 mM PFOA and 14.8 mM PFOS were injected into the macroporous MESIF structure (650 µL per chip). Chips were incubated for 72 h at room temperature to allow methanol evaporation and PFAS adsorption onto the PDMS matrix (Supplementary Fig. S22b). To verify PFAS immobilization, a fluidic setup was used consisting of a PDMS housing with inlet and outlet ports and a central cavity containing PFAS-loaded MESIF (Supplementary Fig. S9a). The system was flushed with 1 mL of water at a flow rate of 10 µL min⁻¹ to assess the stability of PFAS physisorption under flow conditions (Supplementary Fig. S9b). The effluent was collected in PFAS-free HPLC vials (Agilent, USA), and PFAS concentrations were subsequently quantified by HPLC-MS/MS using an Agilent 1290 Infinity II HPLC system coupled to an Agilent 6470 Triple Quadrupole LC/MS detector. Chromatographic separation was performed on an Agilent EclipsePlus C18 RRHD column (1.8 µm, 2.1 × 50 mm) with a Phenomenex C18 guard column, using HPLC-grade water containing 5 mM ammonium acetate and methanol as mobile phases at a flow rate of 0.5 ml min⁻¹.

For environmental cultivation, MESIF chips were filled with 1 mL of the corresponding cultivation medium and used for enrichment experiments In parallel, water samples from the corresponding system (wastewater treatment plant or mesocosm) were consistently collected as controls (hereafter referred to as “Habitat”) In total, nine different treatment conditions were established (Table 1).

**Table 1:** Experimental conditions used in this study.

| Treatment | Medium | Stressor |
| --- | --- | --- |
| M9 | M9 | - |
| M9-G |  | Glyphosate |
| M9-PFOA |  | PFOA |
| M9-PFOS |  | PFOS |
| Native | Native | - |
| N-G |  | Glyphosate |
| N-PFOA |  | PFOA |
| N-PFOS |  | PFOS |
| Habitat | - | - |

### 4.2. Environmental sampling

Two experimental settings were employed: (I) a short-term *in situ* incubation in a wastewater treatment plant (hereafter referred to as “WWTP”), and (II) a long-term laboratory mesocosm incubation (hereafter referred to as “mesocosm”).

#### 4.2.1. Wastewater treatment plant for in situ short-term incubation

MESIF chips were prepared as described above^[30]^. Chips were loaded with either M9 or N medium, resulting in nine treatment conditions (Table 2). For incubation, chips were placed in perforated Falcon tubes (nine openings in the body and five in the cap) to allow water exchange. The tubes were mounted in a cage and suspended within the chemical wastewater tank WP06 B01.1 at KIT Campus North (49.096683, 8.432755). Cultivation experiments were conducted for 21 days, and samples were collected at 9 time points (days 1, 2, 3, 6, 7, 8, 9, 15, and 21) (Fig. 1d).

#### 4.2.2. Mesocosm system for long-term incubation

For long-term mesocosm incubation, a laboratory-scale mesocosm system was built allowing continuous cultivation of MESIF chips for up to 100 days. MESIF chips were prepared and supplemented as described above.^[30]^ Mesocosms consisted of 60 L containers filled with 40 L of wastewater collected from the same wastewater treatment tank used for the *in situ* incubation experiments. Containers were maintained at room temperature under continuous stirring (180 rpm) and covered with plastic foil to minimize evaporation.

Two mesocosms were prepared and operated in parallel, one containing glyphosate-treated MESIF chips and the other PFAS-treated chips; both also included untreated control chips. MESIF chips were placed in permeable mesh bags allowing free water exchange with the surrounding wastewater. Each bag contained nine MESIF chips and three glass beads serving as weights to maintain submersion within the tanks. Samples were collected after 1, 2, 4, 7, 14, 21, 30, 50, 84, and 100 days of cultivation (Fig. 1e).

To assess whether water-soluble glyphosate could rapidly diffuse out of the MESIF into the system, we established small-scale mesocosms adapted to a 10:1 scale (6 L instead of 60 L). Accordingly, we placed only 6 MESIF chips instead of 60 in order to track the release of sodium fluorescein as a fluorescent dye tracer (Supplementary Fig. S1). For this purpose, a 1 mM fluorescein solution was prepared and MESIF chips were loaded with 1 mL per chip. Three chips were supplemented with fluorescein, while three served as controls, enabling monitoring of diffusion dynamics. Samples were collected at the same time points as in the mesocosm experiment over the initial 30 days. Fluorescence in liquid samples was measured using a Synergy plate reader (BioTek, Germany) with excitation at 485 nm and emission at 520 nm. In addition, MESIF matrices were imaged using a gel imaging system (Fusion FX, Vilber, France) to visualize fluorescein distribution within the material. Released fractions in the tank were calculated relative to the initially applied amount.

Finally, to assess physicochemical transformation of PFAS, chips from days 50, 84, and 100 were analyzed. For this purpose, MESIF matrices were extracted from their housing and centrifuged twice at 10,000 × g for 10 min to recover the liquid retained within the material. The collected liquid was subsequently filtered through 0.2 µm filters. The resulting samples were quantified by LC-MS regarding the occurrence shorter-chain PFAS as products of degradation. Due to its lower aqueous solubility, PFOS may exhibit stronger retention within the MESIF matrix, potentially affecting recovery.

### 4.3. Sample processing

#### 4.3.1. DNA Extraction

Genomic DNA (gDNA) was extracted using the DNeasy PowerSoil Pro Kit (QIAGEN, Germany) following the manufacturer’s instructions with minor modifications. For MESIF samples, chips were retrieved from their housings and cut in half. One half was immediately frozen at −20 °C for storage, while the other half was mechanically squeezed in the CD1 lysis buffer provided with the extraction kit to release matrix-associated biomass. Wastewater-derived samples (Habitat) were processed from 2 mL of liquid sample. The samples were centrifuged for 10 min at 10,000 × g, after which the resulting pellet was resuspended in the CD1 buffer and processed according to the extraction instruction.

Mechanical lysis was performed by bead-beating (two cycles of 1 min at 5.0 m s⁻¹; MP Biomedicals, USA). Genomic DNA was eluted in 50 µL nuclease-free water. DNA concentrations were determined spectrophotometrically (NanoDrop OneC, Thermo Fisher Scientific, Germany) and fluorometrically using the Qubit™ dsDNA HS Assay Kit (Thermo Fisher Scientific, Germany). All gDNA extracts were stored at −20 °C until further analysis.

#### 4.3.2. Library Preparation and Sequencing

Metagenomic DNA libraries were prepared using the NEBNext® Ultra II FS DNA Library Prep Kit (New England Biolabs, Germany) according to the manufacturer’s protocol for DNA inputs ≥100 ng. DNA fragmentation was performed for 10 min, followed by library amplification with 12 PCR cycles during the indexing step. Barcoding was carried out using NEBNext® Multiplex Oligos for Illumina (New England Biolabs, Germany). Library concentrations were quantified using the Qubit™ dsDNA HS Assay Kit (Thermo Fisher Scientific, Germany), and fragment size distributions were evaluated using the Agilent High Sensitivity DNA Kit on an Agilent 2100 Bioanalyzer (Agilent Technologies, USA). After normalization to 2 nM, libraries were pooled and sequenced on an Illumina NextSeq 1000 platform (Illumina, USA) using a P2 kit (300 cycles).

### 4.4. Data analysis

#### 4.4.1. Read preprocessing, assembly and coverage estimation

Raw reads were quality-filtered and adapter-trimmed using fastp.^[52]^ Cleaned reads were merged with FLASH^[53]^ and assembled using MEGAHIT^[54]^, generating both individual assemblies and treatment-level co-assemblies. To estimate contig abundance across samples, quality-filtered reads were mapped back to the corresponding assemblies using CoverM^[55]^, and alignment files were processed with samtools.^[56]^ Resulting coverage profiles were used for downstream taxonomic, functional, and genome-resolved analyses.

#### 4.4.2. Community taxonomic and functional profiling

Taxonomic profiling was performed by extracting phylogenetic marker genes from assembled contigs using MDMcleaner^[57]^. Recovered SSU and LSU rRNA genes were taxonomically classified with SINA against the SILVA database.^[58,59]^ Abundance-weighted taxonomic profiles were obtained by mapping reads back to the recovered marker genes using CoverM.

For functional profiling, open reading frames were predicted from assembled contigs using Prodigal^[60]^, and functional annotation was performed with eggNOG-mapper^[61]^ in metagenome mode. Functional annotations were combined with contig coverage values to generate abundance-weighted feature tables, including COG categories^[62]^, KEGG orthologs^[63–65]^, KEGG modules, KEGG pathways, and PFAM domains^[66]^, which were used for downstream analyses of community metabolic potential.

#### 4.4.3. Gene-family clustering, module detection and dark protein analysis

To identify shared and treatment-associated gene families, predicted proteins from co-assemblies were clustered into non-redundant families using MMseqs2^[67]^. Here, and throughout the study, “proteins” refer to predicted open reading frames derived from metagenomic sequence data rather than experimentally validated gene products. Cluster representatives were functionally annotated with eggNOG-mapper, and genes lacking functional annotation were defined as dark proteins. To identify co-occurring gene families, a binary treatment-by-cluster matrix was generated and pairwise Hamming distances were calculated^[68]^. These relationships were visualized using UMAP, and co-occurrence modules, defined as groups of gene families with similar presence/absence patterns across treatments, were inferred using HDBSCAN.^[69,70]^ Modules composed exclusively of clusters lacking eggNOG-based functional annotation were classified as sequence-dark modules. Treatment-associated and sequence-dark modules were further examined, and representative uncharacterized proteins were structurally modeled with AlphaFold2^[71]^ and compared against reference structure databases using Foldseek^[72]^. Finally, proteins that were identified as shared between the short-term WWTP incubation and the long-term mesocosm were further analyzed to predict binding pockets using fpocket^[50]^ and subsequently subjected to docking simulations using gnina.^[51]^ To validate and contextualize the docking results, a control framework was established comprising positive controls (*Homo sapiens* serum albumin (HSA), fatty acid-binding protein 1 (FABP1), and a predicted PFOS-binding reductive dehalogenase^[44]^ and negative controls (triosephosphate isomerase, GAPDH, and Protein A).

#### 4.4.4. Genome-resolved analysis and MAG characterization

Genome binning was performed using a multi-binner strategy combining MaxBin2^[73]^, MetaBAT2^[74]^, and CONCOCT^[75]^, followed by refinement with DAS Tool^[76]^. Binning was applied to both individual assemblies and co-assemblies using coverage-informed approaches. Resulting bins were dereplicated with dRep^[77]^, curated using MDMcleaner^[57]^, and quality-assessed with CheckM2^[78]^. Taxonomic classification of MAGs was performed with GTDB-Tk^[79]^, and phylogenetic placement was inferred using the GTDB-Tk de novo tree workflow. To assess genome-resolved functional potential, genes were predicted with Bakta^[80]^ and annotated using eggNOG-mapper. Genome abundances were estimated by read mapping with CoverM. In addition, genomes of known xenobiotic degraders with available reference sequences were retrieved from NCBI^[81]^ and mapped against the metagenomic datasets to assess their occurrence across treatments.

#### 4.4.5. Data visualization and statistical analysis

All statistical analyses and data visualization were performed using R (RStudio version 2024.12.1; Posit Software, PBC).

### 4.5. Data availability

Metagenomic sequencing reads generated in this study have been submitted to RADAR (DOI 10.35097/fau7wstc5sfgb1xe).

## Acknowledgements

This work was financially supported through the Helmholtz program “Materials Systems Engineering” under the topic “Adaptive and Bioinstructive Materials Systems” (43.33.11), KIT EXU project DigiteLiSE and BMBF project 161L0284A MicroMatrix. The authors thank Simone Weigel and Cornelia Ziegler for their experimental support in sample processing as well as Rafael Peschke and Michael Wagner from Engler-Bunte Institut (EBI), Wasserchemie und Wassertechnologie for conducting the LC-MS measurements.

## Author Information

### Authors

**Marta Velaz Martín** − Institute for Biological Interfaces 1 (IBG-1), Biomolecular Micro- and Nanostructures, Karlsruhe Institute of Technology (KIT), D-76344 Eggenstein-Leopoldshafen, Germany;

**Kersten S. Rabe** − Institute for Biological Interfaces 1 (IBG1), Biomolecular Micro- and Nanostructures, Karlsruhe Institute of Technology (KIT), D-76344 Eggenstein-Leopoldshafen, Germany;

**Laura Meisch** − Institute for Biological Interfaces 1 (IBG-1), Biomolecular Micro- and Nanostructures, Karlsruhe Institute of Technology (KIT), D-76344 Eggenstein-Leopoldshafen, Germany;

**John Vollmers** - Institute for Biological Interfaces 5 (IBG-5), Microbial Genetics and Biotechnology, Karslruhe Institute of Technology (KIT), D-76344 Eggenstein-Leopoldshafen, Germany;

**Anne-Kristin Kaster** - Institute for Biological Interfaces 5 (IBG-5), Microbial Genetics and Biotechnology, Karslruhe Institute of Technology (KIT), D-76344 Eggenstein-Leopoldshafen, Germany;

## Extended Data Figures

**Extended Data Fig. 1.**
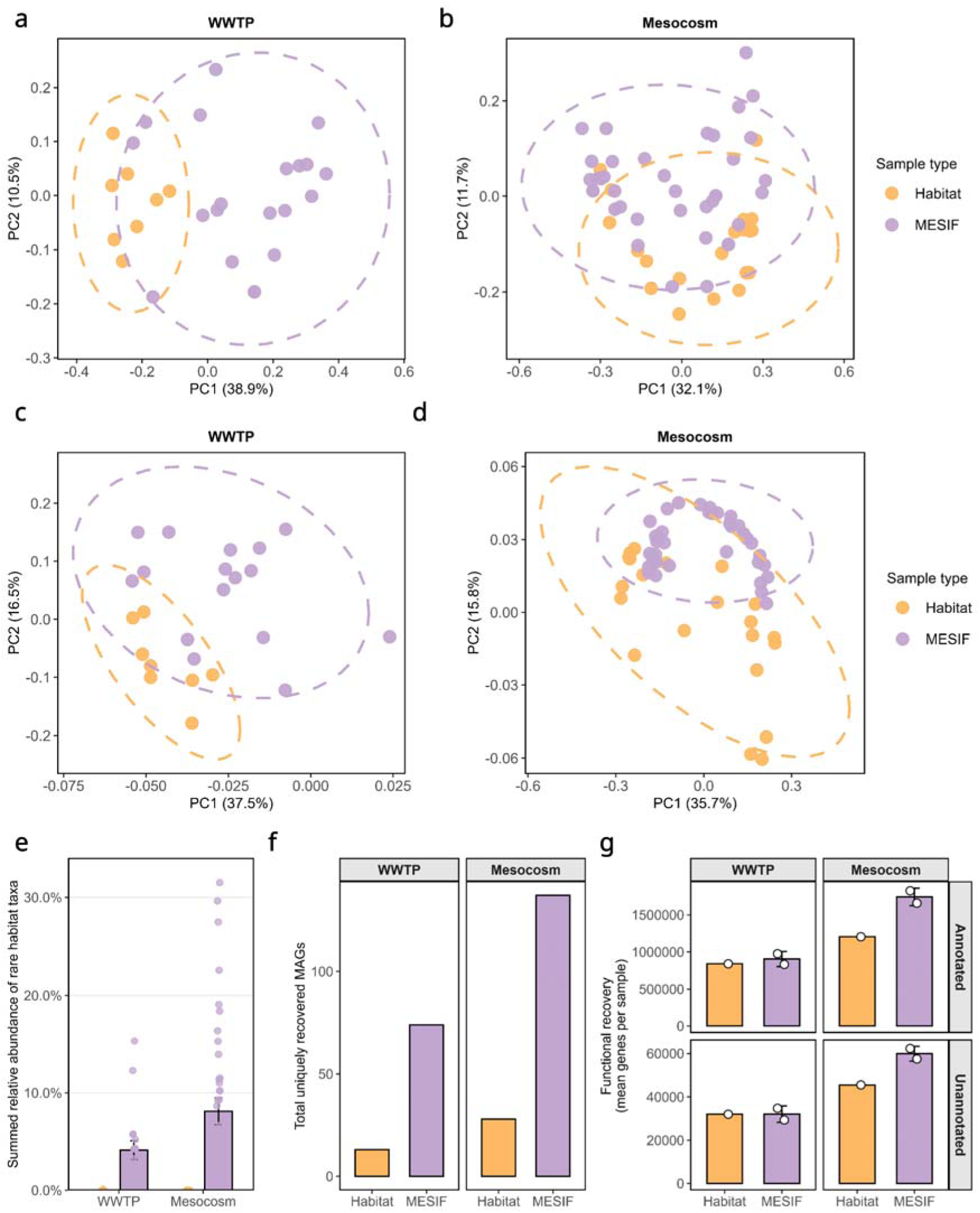
MESIF cultivation drives taxonomic, functional and genome-level divergence from habitat communities. (a,b) Taxonomy-based community dissimilarity between MESIF-associated and habitat-derived microbiomes assessed using Bray-Curtis distances for (a) WWTP and (b) mesocosm samples. (c,d) Functional dissimilarity based on predicted gene content for (c) WWTP and (d) mesocosm samples. (e) Enrichment of low-abundance taxa in MESIF samples relative to habitat communities, indicating selective recruitment of rare populations. (f) Number of reconstructed metagenome-assembled genomes (MAGs) across sample types, reflecting enhanced genome recovery in MESIF compared to habitat conditions. (g) Number of annotated (top) and unannoted genes (bottom) across systems. Center values represent means and error bars indicate standard deviation.

**Extended Data Fig. 2.**
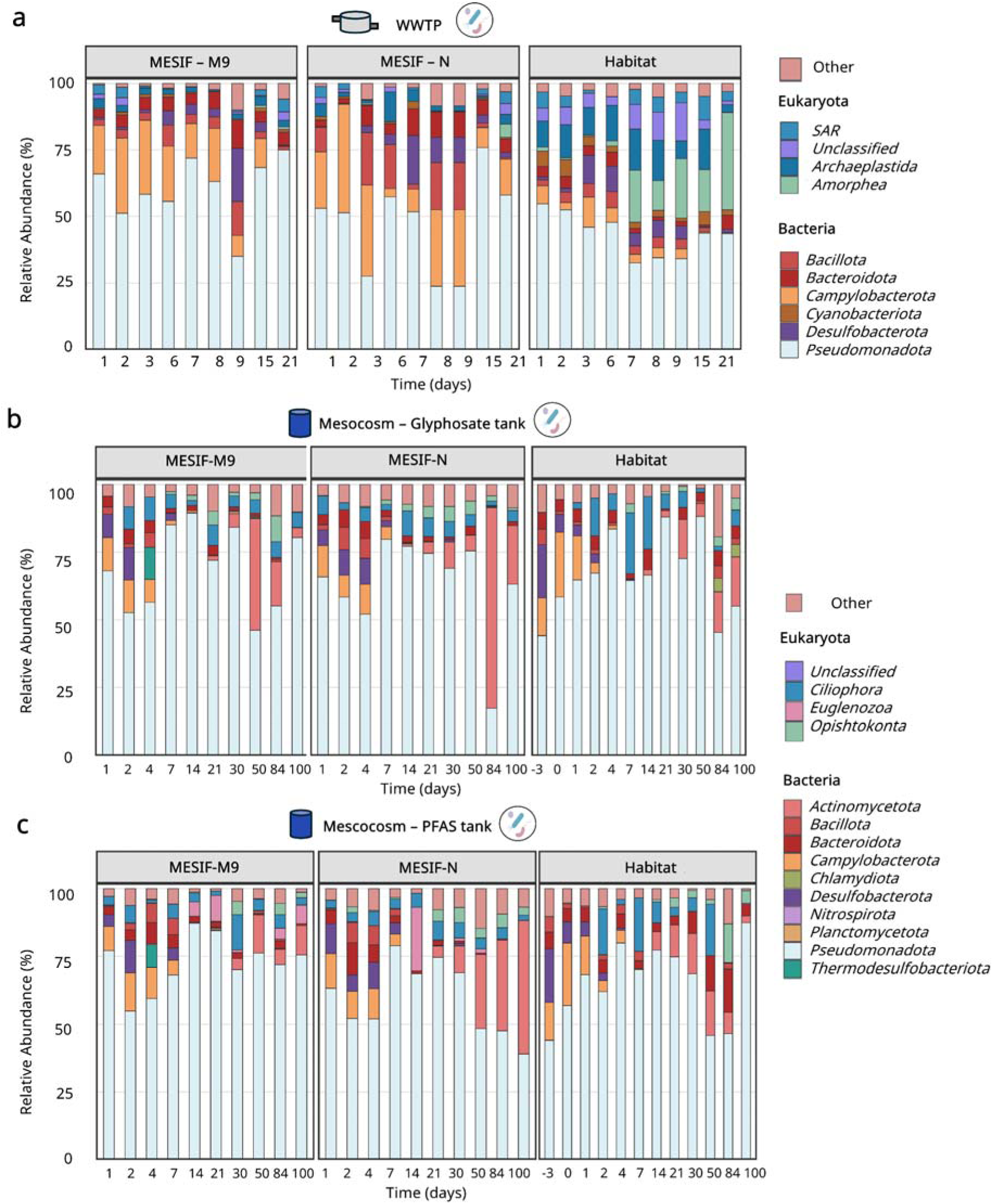
MESIF cultivation restructures microbial community composition in a context-dependent manner across systems and time. (a) Short-term in situ cultivation in a wastewater treatment plant (WWTP). Long-term mesocosm cultivation in the (b) glyphosate-derived or (c) PFAS-derived system, shown here without xenobiotic supplementation to assess system-specific baseline community trajectories; panels (b) and (c) therefore represent independent mesocosm systems and highlight baseline dynamics irrespective of xenobiotic exposure. Stacked bar plots show the relative abundance of microbial phyla over time for control MESIF chips (without xenobiotic supplementation) containing M9 (left) or native wastewater-derived medium (N, middle), compared to the surrounding wastewater (Habitat, right). Each bar represents a sampling time point, and colors indicate the relative abundance of individual phyla. In mesocosm Habitat samples, time point −3 corresponds to the initial WWTP inoculum used to seed the mesocosms, and time point 0 marks the start of sampling following MESIF deployment.

**Extended Data Fig. 3.**
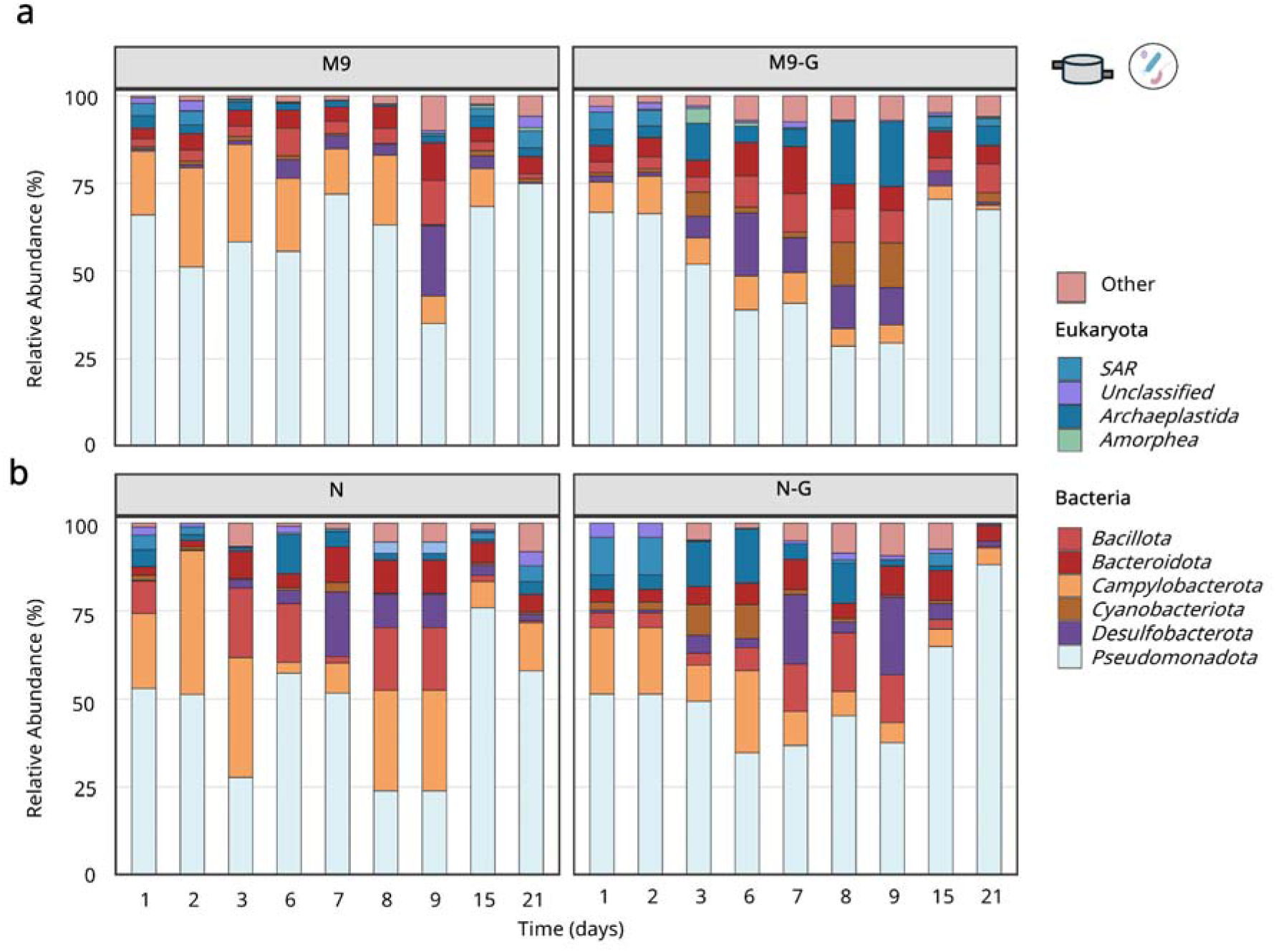
Glyphosate exposure does not induce pronounced taxonomic shifts at the phylum level during short-term WWTP cultivation. Stacked bar plots show the relative abundance of microbial phyla over time for (a) M9 medium comparing control (M9) and glyphosate-supplemented conditions (M9-G), and (b) native wastewater-derived medium comparing control (N) and glyphosate-supplemented conditions (N-G). Each bar represents a sampling time point (days 1-21), and colors indicate the relative abundance of individual phyla. Across both media, community composition remained largely stable at the phylum level, with only minor variation between control and glyphosate-treated samples.

**Extended Data Fig. 4.**
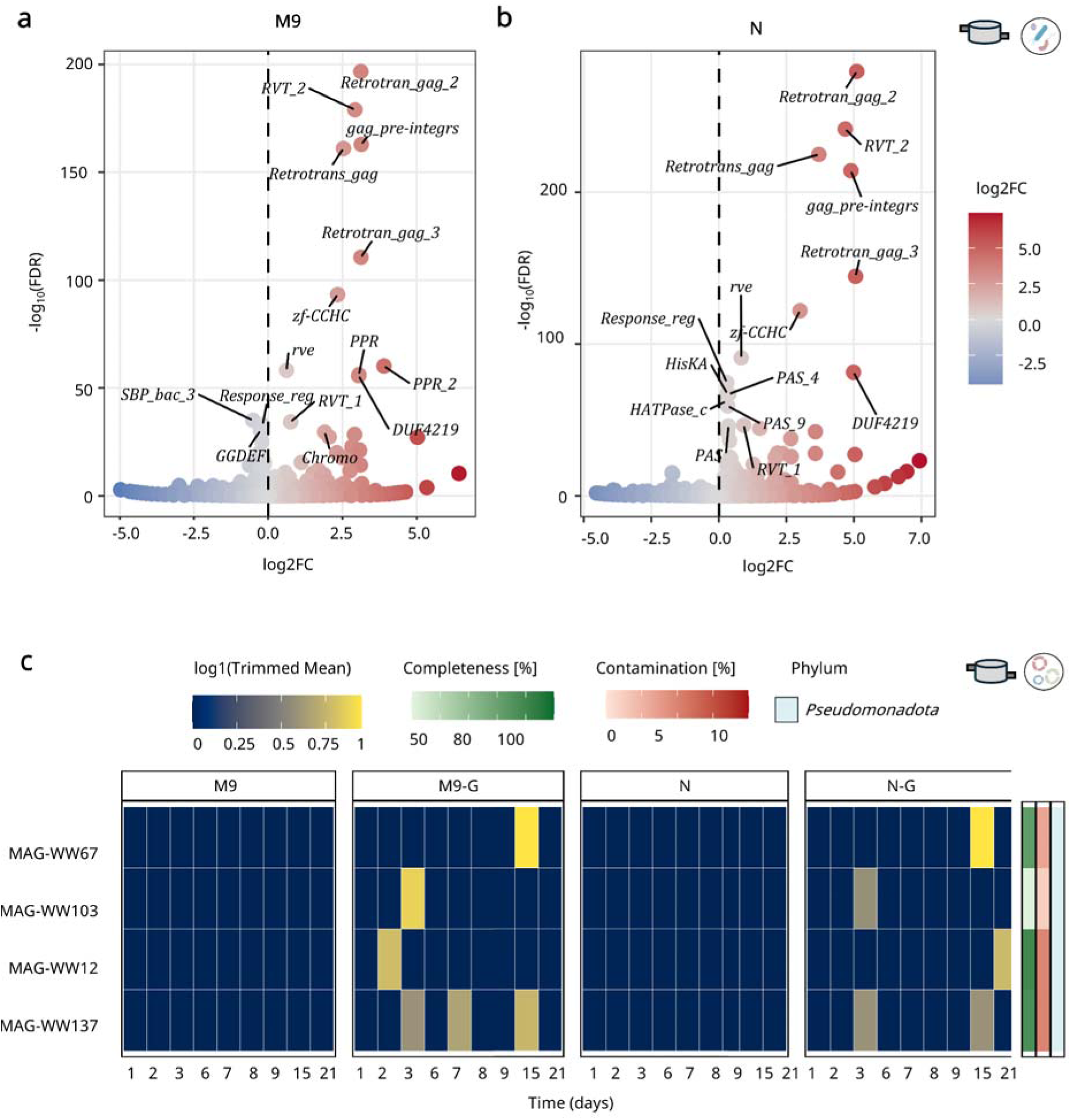
PFAM domain enrichment and genome-resolved dynamics under glyphosate supplementation during short-term WWTP cultivation. (a,b) Volcano plots showing PFAM domain enrichment in glyphosate-supplemented samples relative to controls for (a) M9 and (b) native wastewater-derived medium (N). Each point represents a PFAM domain, with the x axis indicating log_2_ fold change (glyphosate-supplemented vs control) and the y axis representing statistical significance (−log_10_ FDR). Positive values indicate enrichment under glyphosate supplementation, whereas negative values indicate depletion. (c) Genome-resolved analysis of metagenome-assembled genomes (MAGs) uniquely detected in glyphosate-supplemented samples. Heatmaps show the temporal abundance of MAGs across sampling time points, with rows representing individual MAGs and columns representing sampling time points. Values indicate normalized abundance across control (M9, N) and glyphosate-supplemented conditions (M9-G, N-G). Side panels indicate genome completeness, contamination and phylum-level classification.

**Extended Data Fig. 5.**
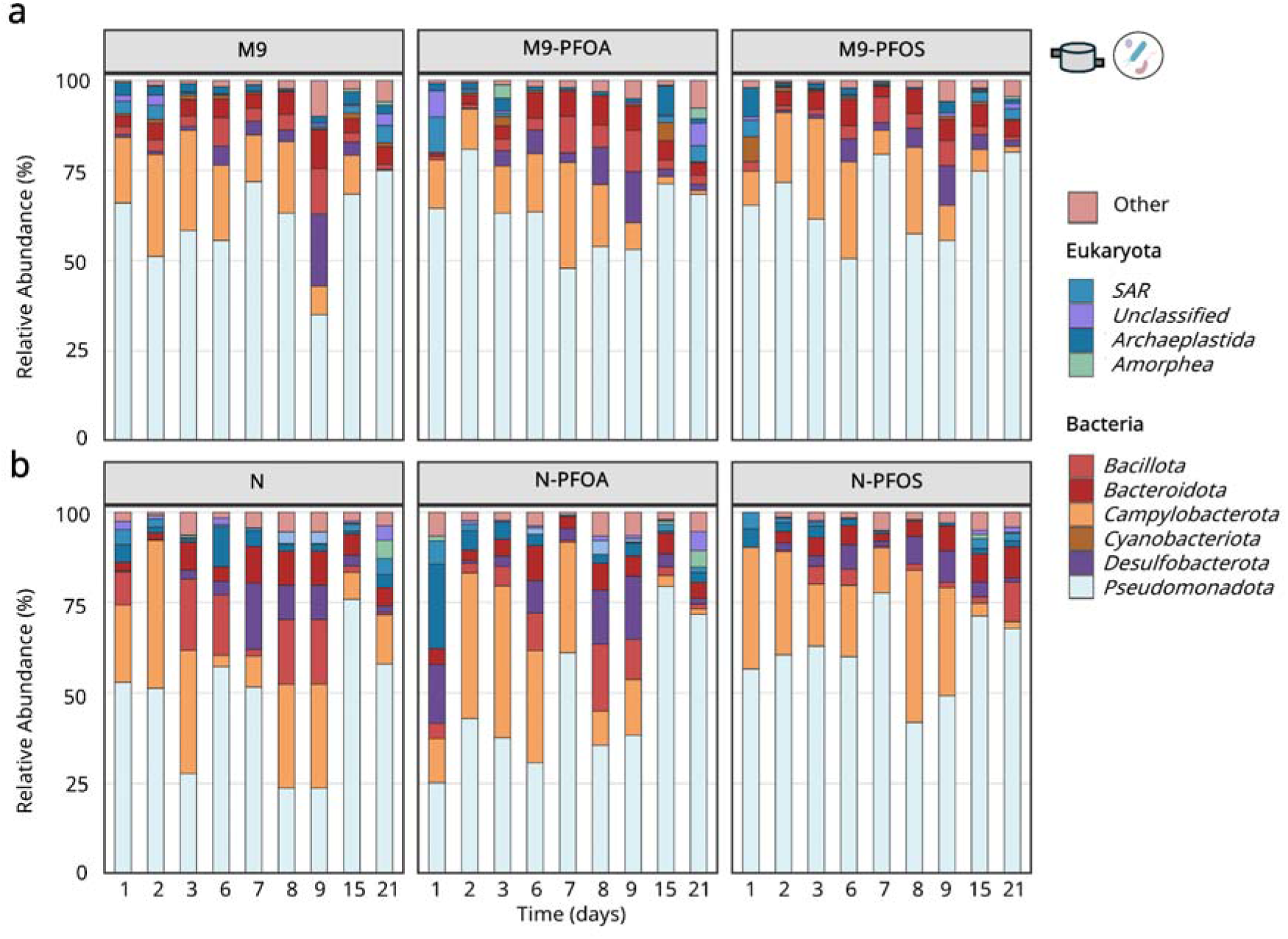
Phylum-level community composition under PFAS exposure during short-term WWTP cultivation. Stacked bar plots showing the temporal distribution of microbial phyla across sampling timepoints for control (left), PFOA-treated (middle), and PFOS-treated conditions (right) in (a) M9 and (b) native (N) medium-supplemented MESIF chips. Each bar represents a sampling timepoint (days 1-21), and colors indicate the relative abundance of individual phyla. Across both media, phylum-level community composition remained largely similar between control and PFAS-exposed samples.

**Extended Data Fig. 6.**
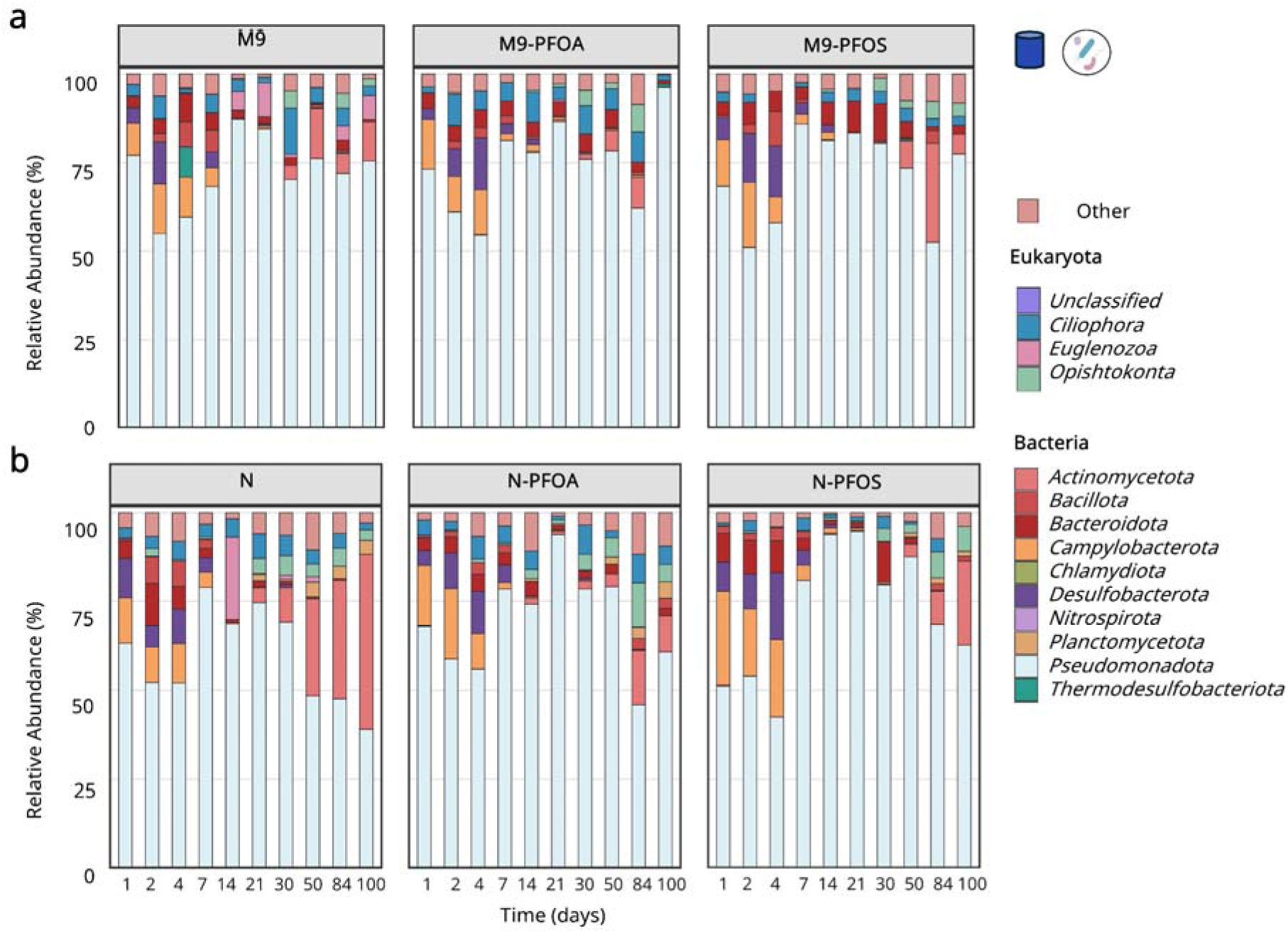
Phylum-level community composition under PFAS exposure during long-term mesocosm cultivation. Stacked bar plots showing the temporal distribution of microbial phyla across sampling timepoints for control (left), PFOA-treated (middle), and PFOS-treated conditions (right) in (a) M9 and (b) native (N) medium-supplemented MESIF chips. Each bar represents a sampling timepoint (days 1-100), and colors indicate the relative abundance of individual phyla. Across both media, phylum-level community composition remained broadly comparable between control and PFAS-exposed samples over time. Although gradual shifts in relative abundance were observed, these were not consistently associated with PFAS treatment.

**Extended Data Fig. 7.**
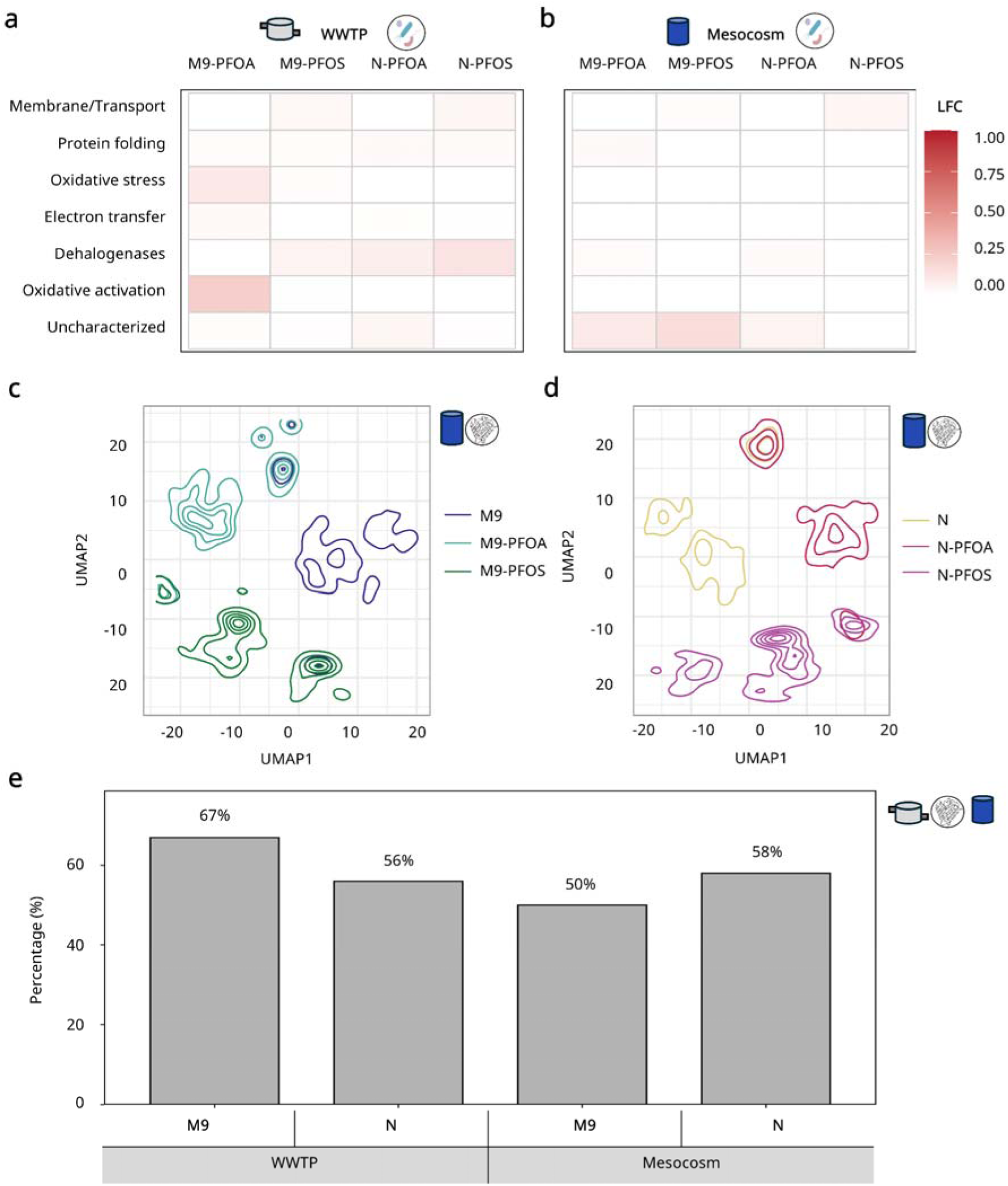
Functional and protein-level responses to PFAS exposure across cultivation systems. (a,b) Functional enrichment of PFAS-associated categories across cultivation systems. Heatmaps show differential abundance (log_₂_ fold change relative to controls) of selected functional categories potentially linked to PFAS-related processes in (a) short-term WWTP cultivation and (b) long-term mesocosm cultivation. Columns represent treatment conditions and rows correspond to functional categories derived from eggNOG-based annotations. Color intensity reflects enrichment relative to the corresponding controls (M9 or N). Across both systems, most categories remained close to baseline levels, indicating limited functional differentiation at the level of annotated pathways. (c,d) Protein-level clustering of mesocosm samples. UMAP-based clustering of proteins after filtering to reduce redundancy, retaining only proteins present in at most n-1 conditions and excluding those shared across all treatments. Density contours illustrate the distribution of protein clusters across control, PFOA-, and PFOS-treated samples for (c) M9 and (d) N communities, highlighting both shared and condition-specific regions of functional space. (e) Structural annotation coverage of PFAS-associated proteins. Bar plot showing the proportion of PFAS-associated proteins that remained uncharacterized after structure-based annotation across experimental systems (WWTP and mesocosm) and media (M9 and N). Despite structural modeling, a substantial fraction of proteins (>50%) remained unannotated, indicating that many sequence-level dark proteins also lack structural annotation.

**Extended Data Fig. 8.**
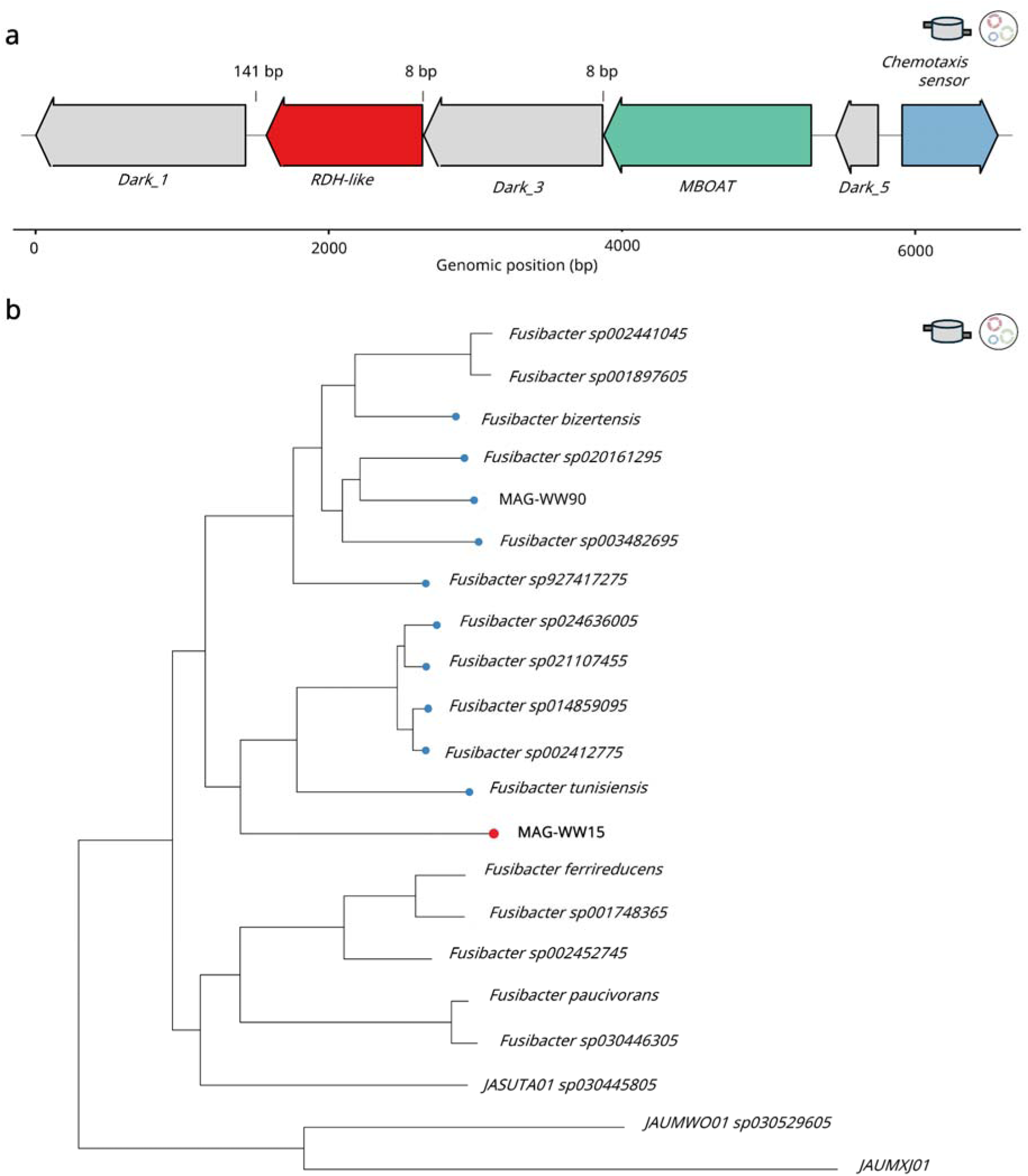
Genomic context and phylogenetic placement of a PFAS-associated MAG encoding a reductive dehalogenase-like protein. (a) Genomic organization of the locus containing a reductive dehalogenase-like (rdh-like) gene (red). The rdh-like gene is flanked by multiple uncharacterized (dark) proteins (grey), a membrane-associated MBOAT protein (green), and a downstream chemotaxis-related sensor (blue), suggesting a functionally linked genomic region potentially involved in environmental sensing or response. (b) Phylogenetic placement of the corresponding MAG (MAG-WW15) based on conserved marker genes. The genome clusters within the genus *Fusibacter*, indicating that it likely represents a previously uncharacterized lineage within this group.

**Extended Data Fig. 9.**
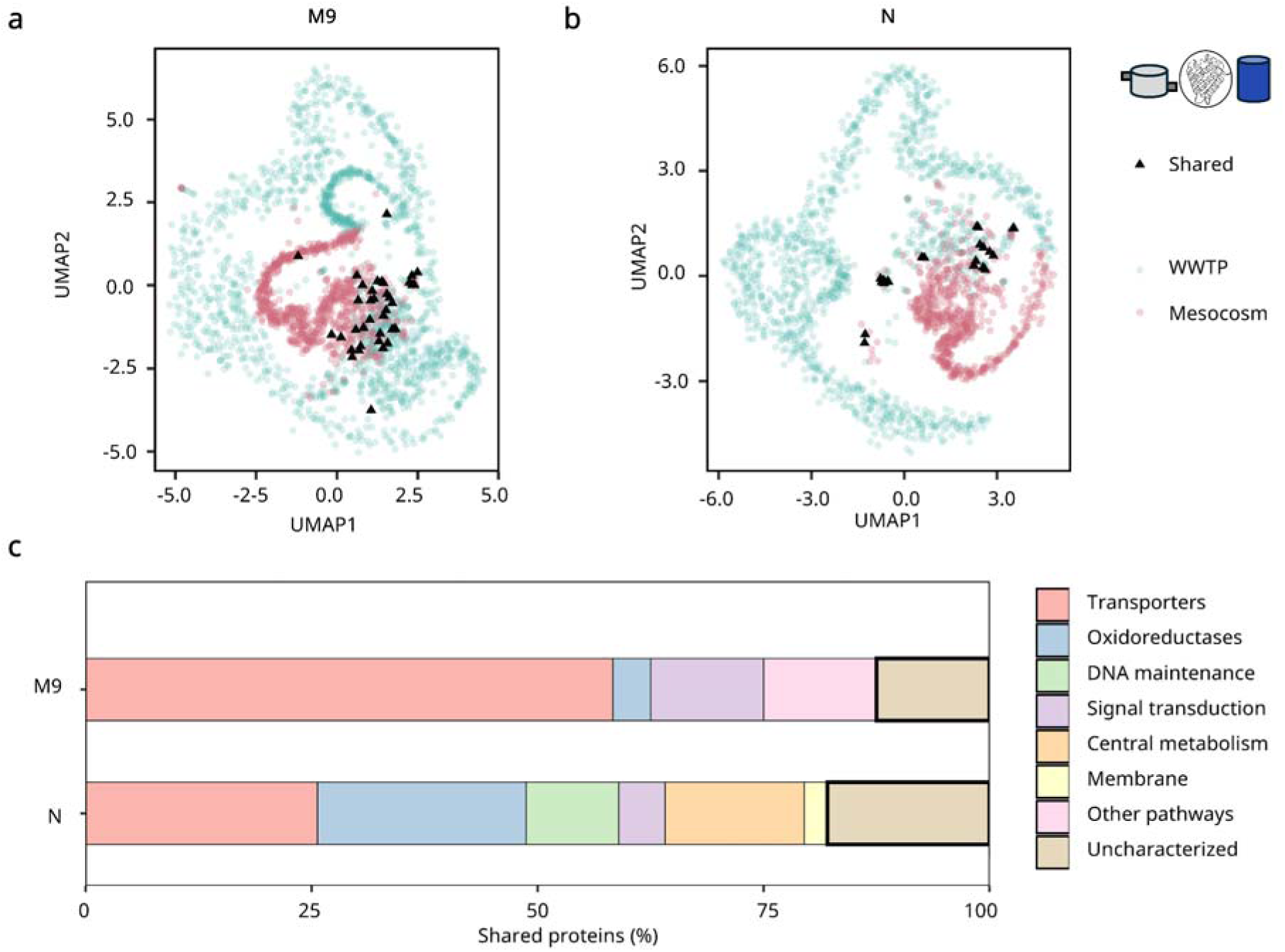
Cross-system comparison reveals limited overlap of PFAS-associated protein space and enrichment of uncharacterized functions. (a,b) UMAP projection of proteins detected across short-term (WWTP) and long-term (mesocosm) cultivation for (a) M9 and (b) native (N) communities. Points represent individual proteins, colored by origin, with proteins detected in both systems highlighted. Limited overlap indicates the absence of a conserved global protein-level signature across systems. (c) Functional classification of proteins detected in both systems, showing a substantial fraction of uncharacterized (“dark”) proteins.

